# Differential gene expression and alternative splicing in insect immune specificity

**DOI:** 10.1101/002709

**Authors:** Carolyn E. Riddell, Juan D. Lobaton Garces, Sally Adams, Seth M. Barribeau, David Twell, Eamonn B. Mallon

**Affiliations:** School of Biological Sciences, University of Edinburgh, U.K.; Department of Biology, University of Leicester, U.K.; School of Life Sciences, University of Warwick, U.K.; Experimental Ecology, Institute of Integrative Biology (IBZ), ETH Zurich, Switzerland

**Keywords:** Genotype-genotype, peritrophic membrane, social insects

## Abstract

Ecological studies routinely show genotype-genotype interactions between insects and their parasites. The mechanisms behind these interactions are not clearly understood. Using the bumblebee *Bombus terrestris* / trypanosome *Crithidia bombi* model system, we have carried out a transcriptome-wide analysis of gene expression and alternative splicing in bees during *C. bombi* infection. We have performed four analyses, 1) comparing gene expression in infected and non-infected bees 24 hours after infection by *Crithidia bombi*, 2) comparing expression at 24 and 48 hours after *C.bombi* infection, 3) searching for differential gene expression associated with the host-parasite genotype-genotype interaction at 24 hours after infection and 4) searching for alternative splicing associated with the host-parasite genotype-genotype interaction at 24 hours post infection. We found a large number of genes differentially regulated related to numerous canonical immune pathways. These genes include receptors, signaling pathways and effectors. We discovered a possible interaction between the peritrophic membrane and the insect immune system in defense against *Crithidia*. Most interestingly we found differential expression and alternative splicing of *Dscam* related transcripts and a novel immunoglobulin related gene *Twitchin* depends on the genotype-genotype interactions of the given bumblebee colony and *Crithidia* strain.

## INTRODUCTION

Invertebrate ecological studies have found infection outcomes within a given host-parasite system are variable. Part of this variance is determined by the interaction of the genotype of the host and the genotype of the parasite [1, 2, 3]. That is this interaction between host and parasite is specific [4]. How is this level of specificity generated? An obvious answer would be an interaction between the parasite and the host’s immune response. We cannot take this for granted however. Various ecological measures of disease outcome have been used to quantify genotype-genotype interactions. These measures include host mortality, fecundity and infection rate. Such measures cannot test directly if it is the immune response that produces this level of specificity [5]. It may be other non-immune processes could explain such outcomes.

The bumblebee, *Bombus terrestris*/trypanosome *Crithidia bombi* system displays host x parasite genotype-genotype interactions [6, 7]. There is evidence that the immune system has a role in generating this host-parasite specific response. A number of studies have found differential immune gene expression in response to *Crithidia* [8, 9, 10, 11]. We found increased *Crithidia* loads in bees whose expression of antimicrobial peptides was knocked down by RNAi [12]. We have even shown that bees from different host genotypes induce differential expression of antimicrobial peptides (AMPs), according to the strain of *C. bombi* they had been infected with [13], that is we found specificity in the immune response itself. A recent paper using RNA-Seq found numerous genes are differentially expressed in a genotype-genotype fashion [14].

Here, we carry out a transcriptome-wide analysis of gene expression in bees during *C.bombi* infection. We have carried out four analyses, comparing 1) expression in infected and non-infected bees 24 hours after infection by *Crithidia bombi* (Infected versus uninfected) 2) expression at 24 and 48 hours after *C.bombi* infection (24 versus 48 hour), 3) searching for differential gene expression associated with the host-parasite genotype-genotype interaction at 24 hours post infection (Specificity) and 4)searching for alternative splicing associated with the host-parasite genotype-genotype interaction at 24 hours post infection (Specificity). Enrichment analysis was also carried out on differential expression data to see which categories of molecules are differentially regulated during infection. The results confirm our previous findings of up-regulation in antimicrobial peptide expression and provide a comprehensive overview of changes in and the specificity of gene expression and alternative splicing after exposure to 2 strains of *C.bombi*.

## RESULTS

The sequences, statistics and annotations for all differentially expressed genes in each of the three differential expression analyses are available in supplementary data (10.6084/m9.figshare.1053093).

### Genes differentially expressed at 24 hours post-infection (Infected versus uninfected)

31,843 unique transcripts were mapped to the transcriptome. 489 transcripts were found to be differentially expressed 24 hours post-infection (FDR <0.05), including 324 downregulated and 165 upregulated transcripts. Reannotating the transcripts using Blast2GO (blastx against the nr database with e <0.001), 109 had no BLAST hits. A further 68 had uninformative BLAST hits (anonymous predicted protein). The remaining 312 were used in the enrichment analysis. Figure 1 shows a summary of the enriched GO terms found (Fisher’s test p <0.05). Defense response (GO:0006952, FDR = 0.047) and chitin metabolism (GO:0006030, FDR = 0.032) were the only processes significantly enriched at a more stringent level (FDR <0.05).

**Figure 1.**
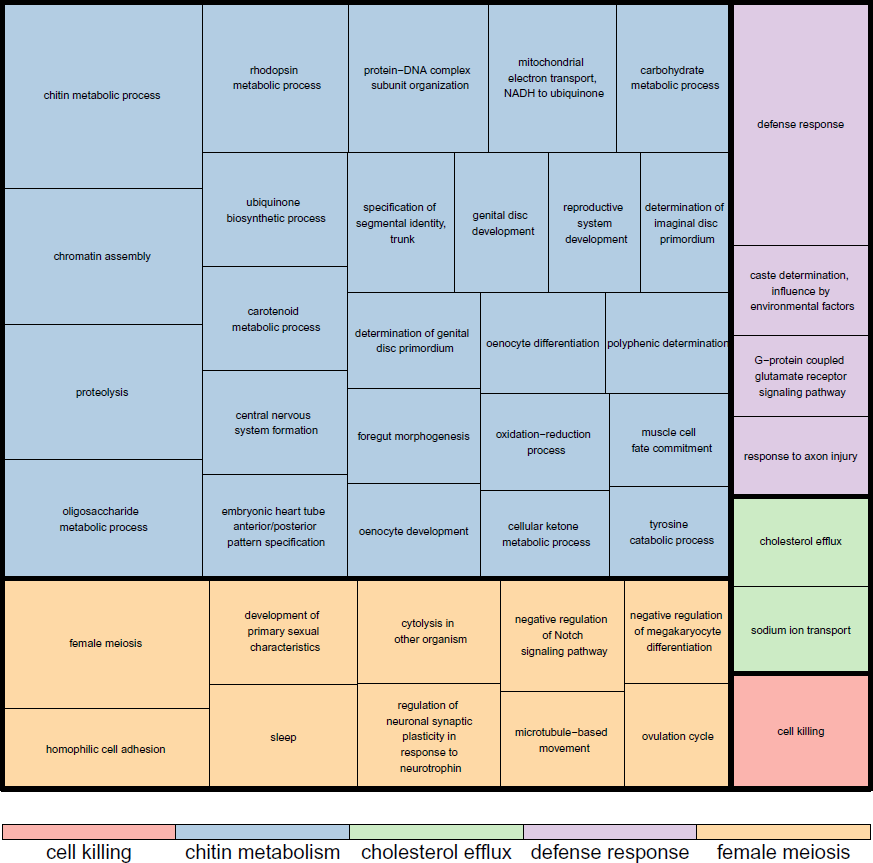
A summary of the enriched GO terms (based on Blast2Go annotation) found for differentially expressed genes at 24 hours post infection (infected versus uninfected). This figure was produced using Revigo

#### Peritrophic membrane

The peritrophic matrix (PM) forms a layer composed of chitin and glycoproteins that lines the insect midgut lumen [15]. The PM facilitates digestion and forms a protective barrier to prevent the invasion of ingested pathogens [16, 15]. *Fibrillin 1* (BTT14121_1), a venom protein precursor (BTT32193_1), *Neurotrypsin* (BTT07956_1), *Peritrophin-1-like* (BTT01709_1, BTT22959_1, BTT37215_1, BTT42262_1) and four chitinase transcripts (*Chitinase 3*: BTT23997_1 BTT38724_1, *Chitinase 4* BTT20684_1, BTT23469_1) are downregulated upon infection. Fibrillins are extracellular matrix macromolecules, ubiquitous in the connective tissues [17]. BTT32193_1 was classed as a venom protein, but was also very similar to *Chitinase 3* (blastx e = 1e^-16^). Chitinases modulate the structure and porosity of the PM [18]. Neurotrypsin is a serine protease expressed in the nervous system [19]. However in the protease domain it shares similarities with Sp22D, a chitin binding serine protease [20]. The chitin fibrils of the PM are assembled into a wide cross-hatched pattern connected by peritrophins [18]. A second group made up of *Peritrophin-1* (BTT05886_1, BTT20661_1) and 3 further chitinase transcripts (*Chitinase 2*: BTT23246_1, *Chitinase 3*: BTT39163_1, *Chitinase 4*: BTT05313_1) is upregulated. Figure 2 shows the correlation of expression patterns between these sixteen transcripts related to chitin metabolism. There is some clustering, but not of any clear functional groups. Taken together this differential expression suggests an important role for the repair or restructuring of the peritrophic matrix in the bumblebees’ response to *Crithidia*.

**Figure 2.**
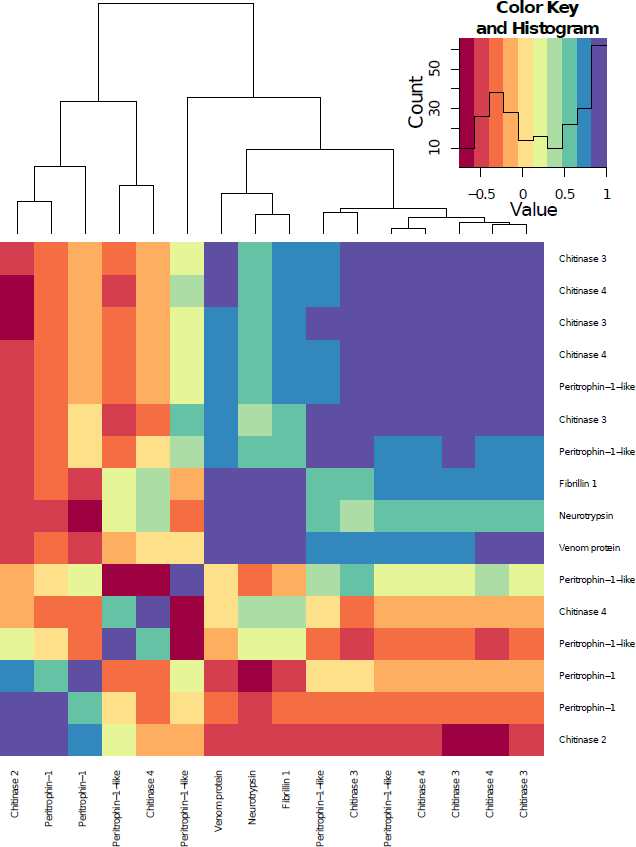
A heatmap showing the correlations of the expression patterns of the transcripts labelled as chitin metabolism genes that where differentially expressed twenty four hours post-infection compared to uninfected samples (infected versus uninfected).

When the BLAST searches against the IIID and nr databases were combined, we found that 89 transcripts relate to canonical insect immune genes. We describe them in the order receptors, serine proteases, signalling pathways and effectors [4].

#### Receptors

The Down syndrome cell adhesion molecule (Dscam), a pattern recognition receptor has come to the forefront of research into insect immune specificity as thousands of different splice forms have been identified and it is associated with insect immunity [21]. We found five downregulated transcripts annotated as immunoglobulin superfamily (*Dscam* included in hit list) (BTT03519_1, BTT08682_1, BTT15814_1, BTT26724_1, BTT27678_1) and one upregulated transcript (BTT03519_1).

#### Serine proteases

Serine proteases are important proteolytic enzymes in many molecular pathways. When these serine proteases are no longer needed, they are inactivated by serine protease inhibitors [22]. CLIP domain serine proteases mediate insect innate immunity [23]. Twenty one transcripts related to serine proteases, serine protease homologues or serine protease inhibitors were differentially expressed upon infection (see Table 1). Lipophorin receptor 2 (downregulated BTT34617_1) binds with serpins to aid in their encytocytosis [24].

**Table 1.** List of transcripts associated with serine proteases and serine protease inhibitors found to be differentially expressed twenty four hours post infection with *Crithidia bombi* (Infected versus uninfected).

| Serine Proteases | Upregulated | Downregulated |
| --- | --- | --- |
| *CLIPA6* |  | BTT20125_1 |
| *CLIPA7* |  | BTT07313_1, BTT31897_1 |
| *CLIPD5* |  | BTT10579_1, BTT10912_1, BTT18247_1 BTT25711_1, BTT06803_1 |
| *SP24* | BTT03436_1 |  |
| *SP27* |  | BTT08108_1, BTT38696_1 |
| *SP35* | BTT05300_1 |  |
| Serine protease homologues | Upregulated | Downregulated |
| *SPH54* | BTT01977_1 | BTT06125_1 |
| Serine protease inhibitors | Upregulated | Downregulated |
| *NEC-like* |  | BTT31997_1 |
| *spn4* |  | BTT04130_1, BTT40693_1, BTT41025_1, BTT41461_1 |
| *SRPN10* |  | BTT04508_1, BTT20259_1 |

#### Signalling pathways

We found a transcript for *Spatzle* (BTT19738_1) downregulated at this time point. Activation of the Toll immune pathway requires the activation of *Spatzle* [25]. MyD88 (upregulated BTT15687_1) is a death domain-containing adaptor activated by Toll leading to the activation of Pelle. *Dorsal* (BTT25273_1) was also downregulated. The nuclear translocation of Dorsal, a member of the NF-kB family, in the Toll pathway induces the expression of many immune genes. We found an upregulated transcript (BTT09662_1) for *Helicase89B* part of the Toll and Imd Pathway. It is required downstream of NF-kB for the activation of AMP genes in *Drosophila melanogaster* [26]. *ird5* (BTT03904_1 downregulated) codes for a catalytic subunit of an IkappaB kinase that cleaves Relish. Relish (Imd pathway) is an essential regulator of antimicrobial peptide gene induction.

In mammals semaphorins are crucially involved in various aspects of the immune response [27]. A *semaphorin-5A-like* transcript (BTT01850_1) was downregulated 24 hours post-infection. Semaphorin regulates the activity of Ras-family small GTPases [27]. A *Ras-like protein11B* transcript (BTT05368_1) was also downregulated. The Ras/MAPK pathway was found to be essential for the suppression of the Imd immune pathway in Drosophila [28].

Drumstick (downregulated BTT13062_1) interacts with the JAK/STAT pathway during its’ development role [29], but we could not find any information about its immune role. Two transcripts (BTT11590_1, BTT14205_1) of *Puckered* were downregulated. *Puckered*, which codes for a dual specificity phosphatase, is a key regulator of the c-Jun-N-terminal kinase (JNK) immune pathway [30]. Mpk2/p38a (downregulated BTT05769_1) is involved in the JNK Pathway and JAK/STAT Pathway. Heat-shock factor activation by p38 is a recently discovered part of antimicrobial reactions in flies [31]. We found two heat shock protein transcripts (BTT23758_2, BTT37030_1) and one other (BTT17701_1) that were downregulated and upregulated respectively. These are all involved in the JAK/STAT pathway.

#### Effectors

In our previous paper [9] we found that antimicrobial peptides were upregulated at 24 hours post-infection. We would expect the same trend here. Indeed, we found that five transcripts for *defensin* (BTT06274_2, BTT8490_1, BTT10405_1, BTT14019_1, and BTT42034_1) and three *hymenoptaecin* transcripts (BTT18071_1, BTT24170_1, BTT24170_2) were all upregulated. An *apidaecin precursor* (BTT33652_1) was downregulated. Apidaecin has recently been shown to be expressed in bumblebees [32] including in response to *Crithidia* [11, 14, 10, 33]. The downregulated beta-amyloid-like protein (BTT20240_1) has been shown to be an antimicrobial peptide in mammals [34]. Hemolectin (BTT15326_1, upregulated) is a clotting protein known to have a role against gram-negative bacteria [35].

Reactive oxygen species (ROS) are generated by respiration in the mitochondria or as part of the immune response [36]. P450 cytochromes are oxidases acting terminally in monooxygenase systems [37]. Some are regulated in response to infection possibly either as direct immune responders [38], producing nitric oxide (NO) or other reactive oxygen radicals or as part of the host detoxification process decreasing oxidative stress after an infection [36]. A number of cytochromes P450 were differentially expressed 24 hours post infection. Ten cytochrome p450 transcripts (*Cyp4p3*: BTT05294_1, BTT20848_1, BTT22253_1, BTT23317_1, BTT32674_1, *cytochrome P450 4g15*: BTT23811_1, BTT32459_1, *cytochrome P450 6k1*: BTT35547_1, BTT40653_1, *cytochrome P450 6a14*: BTT38445_1) were found to be downregulated. Three other cytochrome P450 transcripts (*Cyp4p3*: BTT21216_1, BTT35543_1, *cytochrome P450 315a1*: BTT26726_1) were upregulated. Several other cytochromes (*cytochrome b*: BTT20524_1, BTT39776_1, BTT41896_1, and *cytochrome c*: BTT05255_2) were downregulated.

Numerous other actors in the production of ROS were found to be differentially expressed. *TPX4* (BTT13285_1), coding for a Thioredoxin-dependent peroxidase, which detoxifies hydrogen peroxide was downregulated. This gene was found be differentially expressed during *Plasmodium* infection in *Anopheles gambiae* [39]. *Calcineurin* (BTT08150_1, BTT26273_1) was found to be downregulated 24 hours post-infection, which agrees with our previous findings [9]. In infected *D. melanogaster* larvae, NO signals are enhanced by Calcineurin to promote induction of robust immune responses via the Imd signalling pathway [40].

We found downregulation of *sortilin-related receptor-like* (BTT31654_1). In mammals, sortilin aids in phagocytosis [41]. Two downregulated transcripts (BTT35021_1, BTT08756_1) were matched to *croquemort*, which codes for a key scavenger receptor in the Imd pathway [42]. *Annexin IX* (downregulated BTT02025_1) has been shown to be induced by septic injury in *Drosophila* and is thought to encode for an anticoagulant [43].

#### Miscellaneous

Major royal jelly protein (BTT05317_2, BTT36365_1 upregulated) has been shown to have antimicrobial properties and to be expressed in response to bacterial infection in honeybees [44, 45]. Vitellogenin (downregulated BTT36006_1) is a potent regulator of the immune response in honeybees [46]. Several orthologs of putative *Drosophila* immune loci were differentially expressed 24 hours post-infection (*CG12505*: BTT00934_1, *CG18348*: BTT04397_1, CG7296: BTT15035_1, BTT18395_1, *CG8791*: BTT18908_1, *CG5527*: BTT35653_1, *Fst*: BTT11511_1). The downregulated *CG4393* (BTT05817_1) is weakly analogous to TNF receptor associated factor 3 (TRAF3) that mediates signal transduction involved in mammalian immune responses. Downregulated BTT37289_1 codes for a putative fatty acyl-CoA reductase.

### Genes differentially expressed between 24 hours post-infection and 48 hours post-infection (24 versus 48 hours)

43 transcripts were differentially expressed between 24 hours post-infection and 48 hours post-infection. Of these 17 had no BLAST hits. A further six had uninformative BLAST hits (anonymous predicted protein). The remaining 20 were used in the analysis. Defence response was the only GO term significantly enriched (FDR= 0.00015), with seven transcripts. Three transcripts correspond to *Hymenoptaecin* (BTT18071_1, BTT24170_1, BTT24170_2). They were all upregulated. This suggests a continuing strong AMP production 48 hours after infection. This agrees with other immune assays in bumblebees [47]. Argonaute-2, a RNA-silencing endonuclease, is involved in antiviral defence in insects (downregulated BTT02484_1) [48]. GstD8, a glutathione S-transferase, is involved in the detoxification process (upregulated BTT04810_1) [49]. Dopa decarboxylase (upregulated BTT28048_1) converts L-dopa to dopamine during the melanisation process [50]. SCR-B9 (upregulated BTT40924_1) codes for a scavenger receptor protein. Scavenger receptor proteins have been found to be microbial pattern recognition receptors in flies [51].

### Genes differentially expressed depending on host genotype – parasite genotype interactions (Specificity)

There were 591 differentially expressed transcripts (FDR <0.05). Reannotating the transcripts using Blast2GO (blastx against the nr database with e <0.001), 150 had no BLAST hits. A further 64 had uninformative BLAST hits (anonymous predicted protein). There were 109 transcripts that had previously been found to be differentially expressed at 24 hours post infection.

Of the 591 transcripts, 132 were upregulated and 459 were downregulated. Up or downregulation does not have the same meaning here as in the infected versus uninfected model were there was a clear baseline (uninfected). Our model used colony K strain 8 as the final contrast. From our previously published qPCR data [13], we know the colony K strain 8 interaction displayed the highest levels of AMPs (effectors). Therefore when we say a transcript is upregulated, we mean it is upregulated in this high immune response interaction.

As with the infection data, we combined the BLAST searches against the IIID and nr databases. Ninety transcripts correspond to canonical insect immune genes. We again describe them in the order receptors, serine proteases, signalling pathways and effectors [4].

#### Receptors

Two transcripts were associated with gram-negative binding proteins (upregulated *GNBP*, BTT03533_1 and downregulated *GNBP1-2* BTT35513_1). Although GNBPs are most associated with defense against gram-negative bacteria, they have been show to have a role in response to *Plasmodium* infections [52]. C-type lectins (CTLs) bind carbohydrates and mediate processes including pathogen recognition (Cirimotich et al. 2010). CTL4 is agonist to *Plasmodium* infections in mosquitoes [53]. A *CTL4* transcript (BTT29328_1) was found to be downregulated.

One downregulated transcript was related to *Dscam* (BTT12755_1). A further fourteen downregulated transcripts were part of the Ig superfamily (*IGFn3-1*: BTT05561_1, BTT05581_1, BTT08682_1, BTT12655_1, BTT13442_1, BTT14516_1, BTT18750_1, BTT21156_1, BTT22598_1, BTT22819_1, BTT23339_1, BTT24070_1, *IGFn3-7*: BTT08109_1, BTT09498_1) and one was upregulated (*IGFn3-8*: BTT03519_1). *Dscam* and most of the other Ig superfamily transcripts cluster together in the top right of figure 3, suggesting they are similarly expressed.

**Figure 3.**
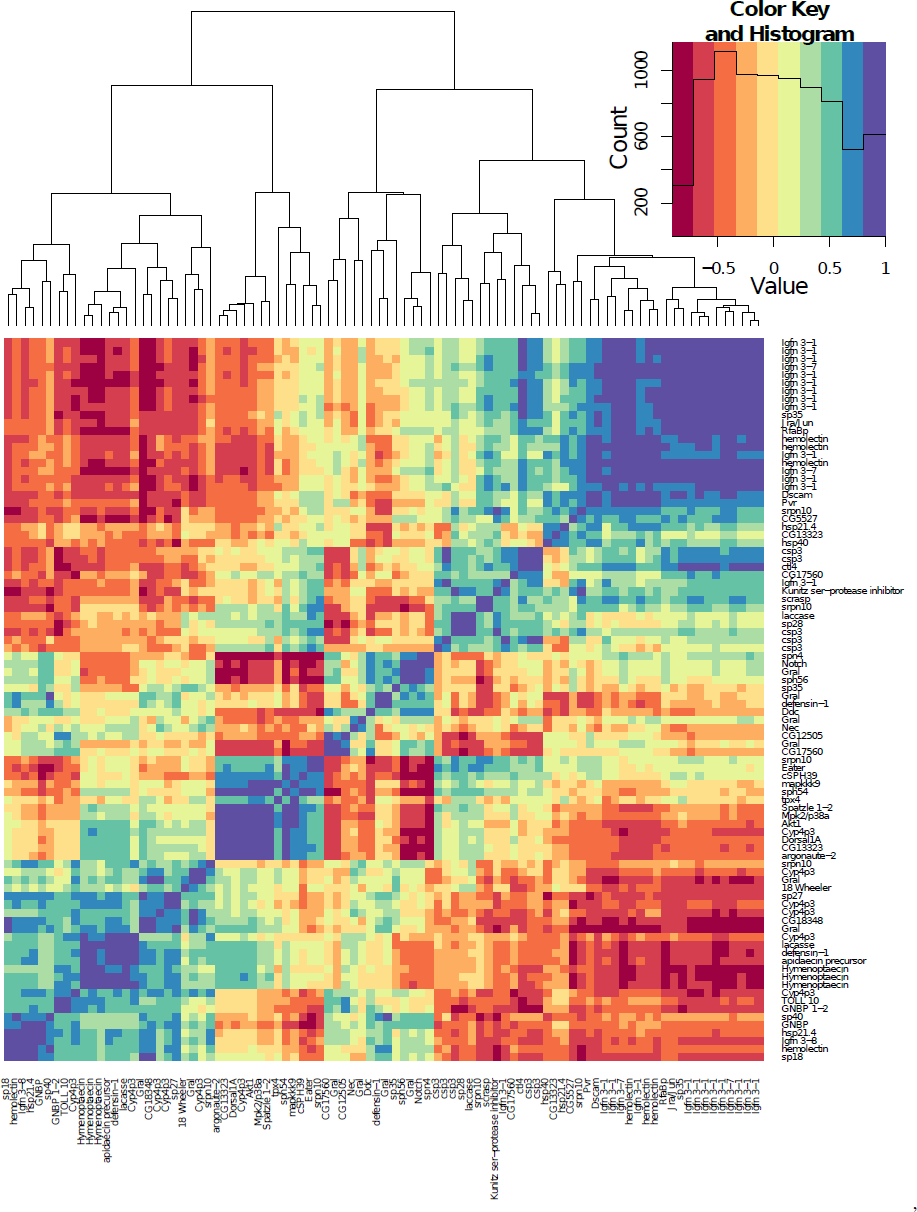
A heatmap showing the correlations of the expression patterns of the 90 transcripts labelled as immune genes in the analysis identifying genes differentially expressed depending on host genotype – parasite genotype interactions (Specifcity).

#### Serine proteases

28 transcripts related to serine proteases, serine protease homologues or serine protease inhibitors were differentially expressed (see Table 2).

**Table 2.**
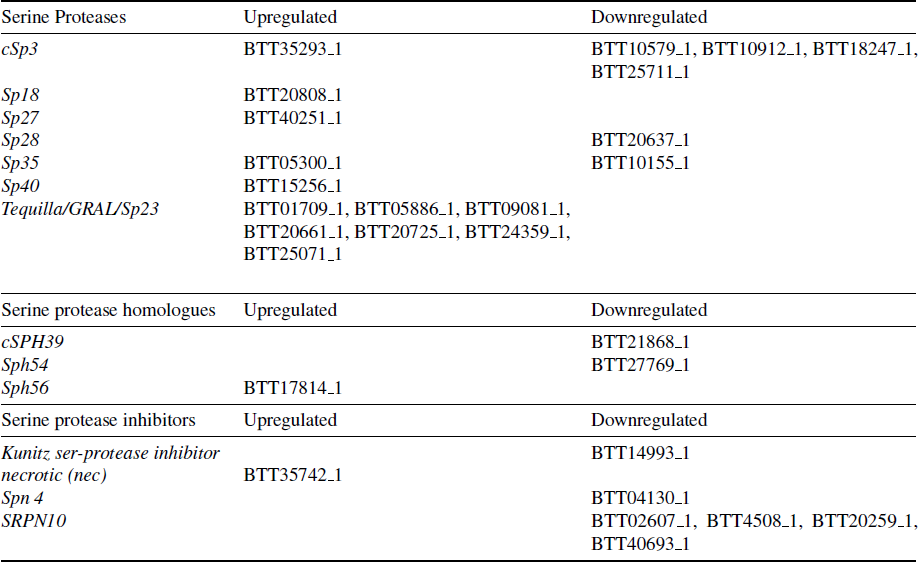
List of transcriptions associated with serine proteases and serine protease inhibitors found to be differentially expressed in the specificity analysis.

#### Signalling pathways

The Toll-like receptor *18Wheeler* (BTT35732_1) and *Toll 10* (BTT09386_1) were both upregulated. 18Wheeler has been shown to be important in the anti gram-negative immune response in Drosophila larvae [54]. *Dorsal 1A* (BTT04010_1), a transcription factor that is an important part of the Toll pathway, was downregulated. A transcript for *Spatzle 1-2* was also downregulated (BTT10679_1).

The tyrosine kinase *Pvr* (BTT04822_1), which inhibits JNK activation [55] was downregulated. *Jun*, a transcription factor of the JNK pathway was downregulated (BTT13636_1). Mpk2/p38a (downregulated BTT16580_1) and MAPKKK9 (downregulated BTT04404_1) are mitogen-activated protein kinases involved in the JNK Pathway and JAK/STAT pathways. We found two heat shock protein transcripts (BTT17371_1, BTT22195_1) and one other (BTT17701_1) that were downregulated and upregulated respectively. These are all involved in the JAK/STAT pathway. Akt 1 (downregulated BTT14188_1) is part of the insulin/insulin-like growth factor 1 signaling (IIS) cascade. IIS plays a critical role in the regulation of innate immunity. Activation of Akt signaling leads to a decrease in malaria infection intensity in mosquitoes [56].

#### Effectors

Five transcripts relate to the AMPs *defensin* (BTT06274_2, BTT42034_1) and *hymenoptaecin* (BTT18071_1, BTT24170_1, BTT24170_2). They were all upregulated. An *apidaecin precursor* (BTT20828_1) was upregulated. *Hemolectin* had three downregulated transcripts (BTT14194_1, BTT17013_1, BTT26614_1) and one upregulated (BTT15326_1). Argonaute-2, a RNA-silencing endonuclease, is involved in antiviral defense in insects (downregulated BTT02374_1) [48].

*Eater* encodes a transmembrane receptor involved in phagocytosis in *Drosophila* (Kocks et al. 2005). A transcript (BTT11132_1) relating to *Eater* was upregulated. The melanisation process component Dopa decarboxylase (BTT19093_1) was upregulated. Another enzyme involved in melanisation, laccase was found to be downregulated (BTT20241_1, BTT33633_1) [57].

*Cyp4p3* transcript BTT40653_1 was upregulated. Two previously unseen *Cyp4p3* transcripts (BTT05254_1, BTT20622_2) were upregulated and one (BTT36257_1) downregulated. *TPX4* (BTT13285_1) that codes for a Thioredoxin-dependent peroxidase was downregulated.

#### Miscellaneous

A small number of transcripts were related to chitin metabolism. SCRASP1 has a chitin-binding domain that has been hypothesized to sense chitin in response to injury and to transduce signals via the serine protease domain [58]. We found an upregulated transcript related to *SCRASP 1* (BTT41923_1). A *peritrophin precursor* was also upregulated (BTT10727_1). As was a *chitinase 3* transcript (BTT23246_1).

*Retinoid and fatty-acid-binding protein (RfaBp)* (BTT07678_1) was downregulated. *RfaBp* was found to be upregulated upon injection of LPS in *Drosophila* during a proteomic study [59]. Notch (upregulated BTT09545_1) is involved in the specification of crystal cells in *Drosophila melanogaster* [60]. Finally, several orthologs of putative *Drosophila* immune loci were found to be differentially expressed (*CG5527*: BTT08512_1, *CG12505*: BTT00934_1, *CG13323*: BTT38025_1, BTT38087_1, *CG17560*: BTT02877_1 downregulated, BTT05845_1 upregulated, *CG18348*: BTT20843_1).

### Genes alternatively spliced depending on host genotype – parasite genotype interactions (Specificity)

The complete output of the DEXSeq analysis is available in the supplemental data (http://dx.doi.org/10.6084/m9.figshare.1053092). 615 loci displayed alternative splicing depending on the the interaction between the host genotype and the parasite genotype. The sequences, statistics and annotations for these loci is available in the supplementary data (http://dx.doi.org/10.6084/m9.figshare.1054540). 98 of the loci had a significant match against the Insect Innate Immunity Database (IIID). Eleven of these are related to receptors including lectins (XLOC_007616 XLOC_007614 XLOC_007613, XLOC_012985, XLOC_011830 XLOC_011825), beta-1,3-glucan recognition protein (XLOC_003146 XLOC_003142 XLOC_003143) and seven transcripts relating to immunoglobulin and fibronectin domains (XLOC_006019, XLOC_006020, XLOC_010845 XLOC_010849, XLOC_012355, XLOC_013957, XLOC_012379, XLOC_014287). Several of these returned Dscam as a blast hit. The transcript with the largest number of alternatively spliced exons was XLOC_010845 XLOC_010849 with 25 variable exons out of 71 total exons (see Figure 4). Against the *Bombus terrestris* genome a blast search of XLOC 010845 XLOC 010849 returned a *twitchin-like* gene (XM_003393691). XLOC_010845 XLOC_010849 aligns with the first 31,160 base pairs of the *twitchin-like* gene (80,078 bp in total). Our cufflinks *Twitchin* gene model matches almost perfectly the one generated automatically by the *Bombus* genome team.

**Figure 4.**
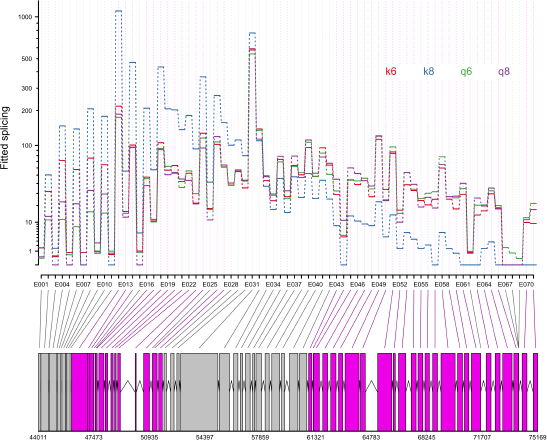
*Twitchin* (XLOC_010845 XLOC_010849) differential exon usage for each of the four host-parasite strain combinations (K6, K8 Q6, Q8). DEXSeq removes the gene-level changes in expression so the differential exon usage is clear. Below is the gene model produced by our cufflinks analysis. The grey blocks are normal exons, the purple blocks represent those exons showing alternative splicing.

## DISCUSSION

We present a comprehensive transcriptomic analysis of gene expression in this important model host-parasite system. We have identified a large number of bumblebee genes whose expression are changed upon infection with *Crithidia*. We also found a large number of genes whose expression depends on the interaction between host and parasite genotypes and therefore show specificity in their expression patterns. A similar number of genes showed alternative splicing in response to parasite-strain interactions.

We confirmed the importance of antimicrobial peptides in the specific defence against *Crithidia* [13, 9, 12]. It is also clear that several other effectors including ROS and phagocytosis may be important. Several immune pathways seem to be important in the anti-*Crithidia* response. These include the Toll, Imd and JAK/STAT pathways. Toll especially seems to be important in a specific immune response.

There are a larger proportion of receptor transcripts showing differential expression found in the specificity analysis (3.2% 19/591) compared to the infection analysis (1.2% 6/489). This is not surprising, as it is be expected that a specific immune response to a given strain would be based mainly on how it is recognised. Although several receptors, including GNBPs and lectins, are differentially expressed, the most exciting discovery is the large number of transcripts related to *Dscam*. The Down syndrome cell adhesion molecule (Dscam), a pattern recognition receptor has come to the forefront of research into insect immune specificity as thousands of different splice forms are generated and it is associated with insect immunity [21]. In the fruit fly *Drosophila*, silencing of Dscam retards the insect’s capacity to engulf bacteria by phagocytosis [61]. In mosquito *Anopheles*, the Dscam splice forms produced in response to parasite exposure differs between bacteria and the malarial causitative agent *Plasmodium* and between *Plasmodium berghei* and *Plasmodium falciparum* [62]. This has been tempered by the finding that Dscam diversity does not increase with exposure to increasing heterogeneity of *Plasmodium falciparum* genotypes [21]. Recently it has been shown that Dscam specificity is mediated by the transcriptional regulation of specific splicing factors downstream of the activation of the Toll and IMD pathways [63]. Our results suggest that Dscam related genes may be important in differentiating strains of the trypanosome *Crithidia bombi*

The alternative splice analysis also found a number of receptor genes. This included numerous *Dscam* related genes. This is encouraging as alternative splicing is the mechanism through which *Dscam* generates the variation that is thought to be useful for immune recognition [21]. The gene with the largest number of alternatively spliced exons was *Twitchin*. This gene was also downregulated 24 hours post-infection (BTT27678_1). Five different transcripts of *Twitchin* (BTT12655_1, BTT13442_1, BTT21156_1, BTT22598_1, BTT23339_1) were expressed in a genotype-genotype fashion in the specificity differential expression analysis. Like *Dscam*, *Twitchin* possesses a large nymber of fibronectin and immunoglobulin domains. *Twitchin* is part of the *titin* family of genes. They produce large filamentous proteins that mediate the transduction of mechanical signals in muscles [64]. However a *titin* gene was found to show differential exon usage depending on if a mosquito was infected with bacteria or *Plasmodium* [65]. *Twitchin* is an exciting possible candidate gene for the source of specifcity in this system and will be the subject of future work in our lab.

We found a number of genes associated with chitin metabolism differentially regulated 24 hours post-infection. Through several pathways chitin metabolism is fundamental to invertebrate immunity [66]. As an aside, an intriguing hypothesis is that chitin metabolism is the nexus through which defense against predators and against parasites are traded-off [66]. Our data suggests that the peritrophic matrix may be fundamental in the bee’s defence against *Crithidia*. The peritrophic matrix acts as an immunological barrier against trypanosomes. Tsetse flies with an underdeveloped PM have lower levels of refractoriness to trypanosome infections [67]. This is due to a premature immune response; the trypanosomes get through the PM quicker and stimulate the immune response at an earlier stage compared to refractory flies.

A recently published paper by one of the authors, found genotype x genotype interactions in the expression of a smaller number of genes [14]. We hypothesise that our much larger catalogue of genes, including *Dscam* and *Twitchin*, is due to our experimental design. Our samples were preselected based on that they showed extreme reciporcal specificity in AMP expression [13]. The two colonies used were from different populations, one wild caught and one commercial. This increased the potential differences in their response to the two strains. In turn, this increased our likelihood of detecting differential expression and exon usage using RNA-seq.

In this paper we have shown that the expression and alternative splicing of immune genes is associated with specific interactions between different host and parsite genotypes in this bumblebee / trypanosome model. In future RNAi work we will knockdown candidate genes thereby altering these specific interactions to directly examine their biological significance.

## METHODS

The samples used during this experiment are as previously described [13]. We have chosen samples that displayed a reciprocal pattern of expression for the three antimicrobial peptides (AMPs) tested in that paper. These were colony K5 (called K from now on) and Q1 (Q) and strains 6 and 8. K-8 showed a high AMP expression, Q-8 a low expression level, Q-6 a high level and K-6 a low level of AMP expression.

### Sample collection

Experiments were carried out on one commercially reared bumblebee colony from Koppert Biological Systems U.K. (Colony K) and one colony from a wild caught queen (Colony Q). Faecal samples from these colonies were checked under a light microscope to ensure there was no *Crithidia bombi* present [68]. All parasite isolates used originated from wild queens collected in Spring 2008 in the University of Leicester botanic garden. The *Crithidia* from each individual queen was infected into a group of 10 workers from a different colony to amplify the strain and to provide a source for experimental infections. Experiments began when the colonies had a minimum of thirty workers, approximately four weeks old. Between observations, colonies were fed *ad libitum* with pollen (Percie du sert, France) and 50% diluted glucose/fructose mix (Meliose – Roquette, France). Before and during the experiments colonies were kept at 26°C and 60% humidity in constant red light.

### Infections

To prepare *C. bombi* isolates, faeces was collected from infected workers and mixed with 50% diluted Meliose to create a standardized dose of 500 *Crithidia* cells per microlitre of inoculum. Previous studies had shown that such inocula, prepared from different colonies, are genotypically different [7] and generate specific responses in novel hosts [6]. We chose this method over *in vitro* culturing to prevent possible attentuation of strains’ infectivity associated with culturing [69]. One possibility is that by using faeces we may be introducing hidden infections or gut microbiota from the donor queens. We have attempted to mitigate this by using bees from a single colony to culture and grow the *Crithidia*. Although the queens faeces may indeed contain hidden infections and microbiota, they all must be passed through the same host background before they are used experimentally. We infected a sample of workers from each of K and Q bumblebee colonies (representing different host lines) with an inoculum of faeces from each of the two wild infected queens (6 and 8 *Crithidia* strain). We also collected uninfected controls, which were sacrificied at five days old (fours days plus 24 hours). Bees were four days old at the time of infection. Bees were collected over several days and distributed across treatment groups [70]. After infection bees were kept in colony x strain groups (1–3 individuals depending on day collected) and fed *ad libitum*. Twenty four hours or 48 hours post infection the bees were sacrificed by freezing in liquid nitrogen and stored at minus 80 °C.

### RNA sample preparation and sequencing

Total RNA was extracted from 23 individual homogenised abdomens using Tri-reagent (Sigma-Aldrich, UK). Samples (Colony-Strain-Timepoint (number of replicates)) were K-6-24 (3), K-6-48 (3), K-8-24 (3), K-8-48 (3), K-Uninfected (2), Q-6-24 (3), Q-6-48 (3), Q-8-24 (2), Q-uninfected (1). Any residual contaminants were removed from the RNA using the RNeasy mini kit (Qiagen, UK) and manufacturer’s RNA clean-up protocol. To remove residual genomic DNA, RNA samples were treated with DNase (Sigma-Aldrich, UK). TruSeq RNA-seq libraries were made from the 23 samples at NBAF Edinburgh. Sequencing was performed on an Illumina HiSeq ® 2000 instrument (Illumina, Inc.) by the manufacturer’s protocol. Multiplexed 50 base single-read runs were carried out yielding an average of 12M reads per sample.

### Differential gene expression analysis

The reference transcriptome was downloaded from http://www.nematodes.org/downloads/databases/Bombus_terrestris/ [32]. Functional annotation related to the transcriptome was obtained using the BLAST2GO package [71]. Alignment was done using GSNAP (version 2012-07-20) [72]. Only reads that mapped uniquely were selected for further analysis. Counts were generated per transcript for each sample.

Differential expression analysis was performed using the edgeR (3.4.0) package [73] in R (3.0.1) [74]. Normalization factors were computed using the TMM technique, after which tagwise dispersions were calculated and subjected to a generalized linear model (GLM). Resulting p values were subjected to Benjamini–Hochberg multiple testing correction to derive FDRs; only transcripts with a FDR <0.05 were considered for further analysis. Three separate GLMs were carried out. One looked for transcripts that are differentially expressed upon infection with *Crithidia* at 24 hours post-infection (Infected versus uninfected) (0+colony+infect(yes/no)). “Infect” here are bees infected with either strain 6 or 8. Another GLM looked at the gene expression difference between 24 hours and 48 hours post strain 6 infection (24 versus 48 hours)(0+colony + time). The third GLM looked for transcripts that were expressed in a specific pattern at 24 hours post-infection (specifcity)(0+colony*strain).

Using Blast2Go, we then carried out an enrichment analysis (Fisher exact test) on each of these lists of differentially expressed genes to see which GO terms are overrepresented relative to the entire genome. We then used REVIGO to summarize and visualise these terms [75].

For each of the lists of differentially expressed transcripts we also carried out a blastx analysis against the insect innate immunity database (IIID) [76]. We used the BLOSUM62 matrix with a word size of 3. The results were filtered to only contain hits with an E-value <1e^-10^ and a bit score ≥30.

### Alternative splice analysis

The eleven samples used in the specificity analysis above were also tested for alternative splicing.

#### Alignment and creation of gene set

Reads were first aligned to the *Bombus terrestris* reference genome (AELG00000000.1) using the fast splice junction mapper Tophat [77]. Preliminary sequence data was obtained from Baylor College of Medicine Human Genome Sequencing Center website at http://www.hgsc.bcm.tmc.edu. The Tophat produced alignment files were then passed to Cufflinks to generate a transcriptome assembly for each sample [78]. These assemblies were then merged together using the Cuffmerge utility [78], into a general transcriptome assembly.

#### DEXSEQ analysis

The BAM files from the Tophat analysis were converted into SAM format using SAMTools [79]. The GTF file from Cuffmerge was flattened into a GFF file with collapsed exon counting bins using the Python script dexseq_prepare_annotation.py found in HTSeq [80]. For each SAM file, python_count.py (HTSeq) counts the number of reads that overlap with each of the exon counting bins defined in the flattened GFF file. We tested for differential exon usage using the R package DEXSeq [81]. We used a full model of counts = sample+exon+(colony+strain)*exon+(colony:strain)*I(exon==exonID) and a reduced model of counts = sample+exon+(colony+strain)*exon. This identified genes which show differential exon usage due to the interaction of strain and colony (FDR <0.01).

#### Blast analysis

We extracted the nucleotide sequence for all differentially expressed transcripts and searched for any matching sequence on NCBI using BLASTn [82] with an E-value cutoff of 0.001, restricting the sequences to those from *B. terrestris*.

## ACKNOWLEDGEMENTS

Thanks to Simon Anders and Alejandro Reyes for help with the DEXSeq model Thanks to Paul Schmid-Hempel for discussions. CR was funded by a BBSRC studentship. This work was partially funded through a NERC NBAF pilot grant (NBAF606) to EBM. We thank the Bumblebee Genome Consortium (http://hymenopteragenome.org/beebase/) for providing genomic resources that were used for this study.

